# iSeq: An integrated tool to fetch public sequencing data

**DOI:** 10.1101/2024.05.16.594538

**Authors:** Haoyu Chao, Zhuojin Li, Dijun Chen, Ming Chen

## Abstract

High-throughput sequencing technologies (Next Generation Sequencing; NGS) are increasingly utilized by researchers to tackle a diverse array of biological inquiries. Leveraging the remarkable scale and efficiency of modern sequencing, significant advancements are made across various fields, spanning from genome analysis to the intricate dynamics of protein-nucleic acid interactions. Recognizing that NGS data harbors rich biological information, the International Nucleotide Sequence Database Collaboration (INSDC) was established nearly 40 years ago to collect and disseminate public nucleotide sequence data and associated metadata. The National Genomics Data Center (NGDC) has also provided open access to vast amounts of raw sequence data. These databases have greatly enhanced the capacity for reanalyzing NGS data. In recent years, amid the rise of large language models, biological sequences and data have emerged as inputs for training models to address biological challenges. However, methods for programmatically accessing this public sequencing data remain limited. To address this gap, we have developed iSeq, an integrated tool that allows for quick and straightforward retrieval of metadata and NGS data via the command-line interface. iSeq is currently the only tool that supports simultaneous retrieval from multiple databases (GSA, SRA, ENA, DDBJ, and GEO). Additionally, iSeq supports a wide range of accession formats as input and features parallel downloads, multi-threaded processes, and FASTQ file merging. It is freely available on Bioconda (https://anaconda.org/bioconda/iseq) and GitHub (https://github.com/BioOmics/iSeq).

**Highlights:**

- iSeq supports multiple databases for accessing a wide range of raw sequencing data and metadata.
- iSeq supports at least 25 different accession formats as input.
- iSeq supports parallel downloads, multi-threaded processes, FASTQ file merging, and integrity verification.

## INTRODUCTION

Next Generation Sequencing (NGS) represents a significant advancement in molecular biology, allowing for high-throughput sequencing of DNA and RNA [1]. Alongside the development of NGS technology, numerous omics applications have emerged, including bulk RNA-Seq [2], ATAC-Seq [3], ChIP-Seq [4] Hi-C [5], and WGS [6]. Exciting new applications are also being explored, such as single-cell and spatial omics in epigenomics [7], transcriptomics [8], proteomics [9], and metabolomics [10]. These technologies have revolutionized various fields. For instance, in oncology, NGS is extensively used to identify genetic mutations in tumors, aiding in the development of targeted therapies [11]. Moreover, during the COVID-19 pandemic, NGS played a crucial role in identifying and tracking the SARS-CoV-2 virus [12]. Beyond medical applications, NGS is utilized in agriculture to improve yields, quality and resistance by studying plant genomes [13].

With the widespread adoption of NGS technology, the volume of NGS data has dramatically increased. Large-scale mining and analysis of NGS data can provide a more comprehensive understanding of biological mysteries. For example, Gavish *et al*. utilized 1,456 public single-cell RNA-seq (scRNA-seq) samples to uncover hallmarks of transcriptional intratumour heterogeneity across a thousand tumors [14]. Zuntini *et al*. built the tree of life for almost 8,000 angiosperm genera using the publicly available multi-omics data of 2,054 species and their internal NGS data [15]. Nowadays, in the era of flourishing large language models, large-scale NGS data can significantly enhance model performance. For instance, Genoformer used 561 publicly available scRNA-seq datasets (about 30 million cells) to train a model, accelerating the discovery of key network regulators and candidate therapeutic targets [16]. Similarly, scGPT used 65 scRNA-seq datasets (over 33 million cells) to train a model optimized for superior performance across diverse downstream applications, such as cell type annotation and multi-omic integration [17].

New NGS data are increasingly generated from economically growing countries, and data reuse and reanalysis are becoming common [18]. To support NGS data storage and sharing for the global scientific community, the International Nucleotide Sequence Database Collaboration (INSDC) was established nearly 40 years ago to collect and disseminate public nucleotide sequence data and associated metadata [18]. INSDC includes three partner organizations: the DNA Data Bank of Japan (DDBJ; http://www.ddbj.nig.ac.jp) in Japan [19], the European Nucleotide Archive (ENA; https://www.ebi.ac.uk/ena) in the UK [20], and GenBank (https://www.ncbi.nlm.nih.gov/genbank) in the USA [21]. The Sequence Read Archive (SRA; https://www.ncbi.nlm.nih.gov/sra) was established as part of INSDC in 2009 [22]. In addition to INSDC, the Genome Sequence Archive (GSA; https://ngdc.cncb.ac.cn/gsa) at The National Genomics Data Center (NGDC) in China also provides vast amounts of raw sequence reads [23]. These organizations collaboratively maintain NGS data for the benefit of science and the wider community.

While high-throughput DNA sequencing technologies have revolutionized biological research, the continuously growing volume of data presents challenges for efficient and convenient access and deep mining. In this context, we introduce iSeq, a tool designed to streamline the retrieval of metadata and NGS data from GSA, SRA, ENA, and DDBJ databases. iSeq aims to facilitate efficient data access, supporting multiple accession formats and offering functionalities such as parallel downloads and multi-threaded processes.

## METHODS

iSeq is a comprehensive Bash script designed to efficiently fetch public sequencing data from multiple bioinformatics databases, including GSA, SRA, ENA, and DDBJ. The workflow begins by providing an accession number **(Figure 1)**, which can be in various formats such as Project, Study, Sample, Experiment, or Run. iSeq automatically detects the accession format and fetches metadata from the appropriate source, prioritizing ENA among the partner organizations of INSDC or GSA due to their extensive data availability.

**Figure 1.**
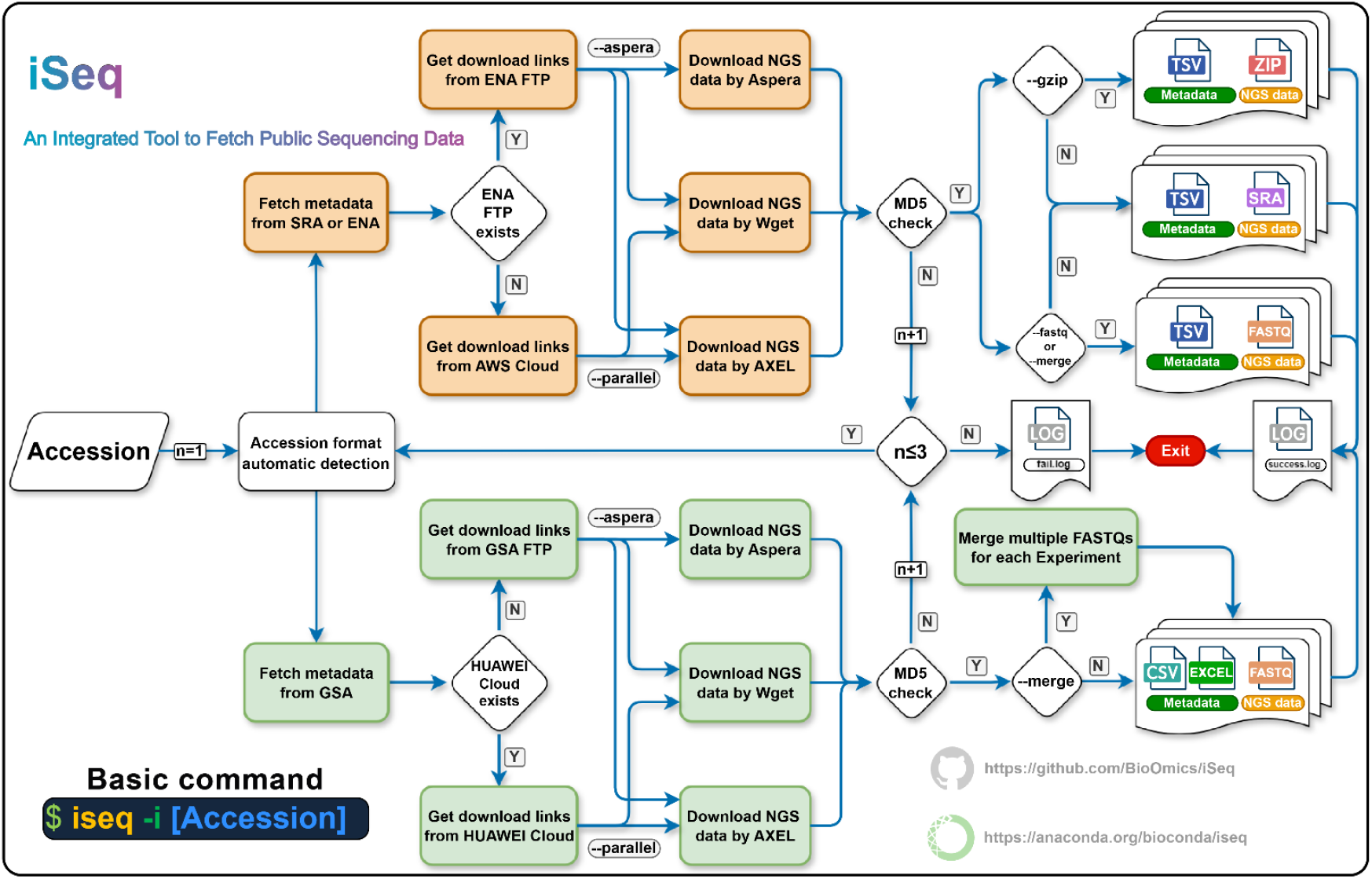
iSeq workflow pipeline. Once iSeq receives the correct accession, it will execute from left to right in sequence.

Once the metadata is fetched, iSeq determines the best source for downloading the sequencing data. If the fetched accession is in the INSDC databases and the FTP channel is available in ENA, iSeq will proceed to download the data using either Aspera (https://www.ibm.com/products/aspera) for fast transfer, Wget (https://www.gnu.org/software/wget), or AXEL (https://github.com/axel-download-accelerator/axel) for parallel downloading. In cases where the FTP channel is not available in ENA, iSeq will fetch download links from AWS Cloud channel, again utilizing Wget or AXEL based on user preferences. If data is stored in GSA and the HUAWEI Cloud channel is accessible, iSeq prioritizes accessing the HUAWEI Cloud channel for its speed and reliability. Alternatively, iSeq will use the FTP channel with Aspera, Wget, or AXEL for downloading. After the initial download, iSeq will conduct an MD5 checksum to ensure the integrity of the downloaded files. If the checksum does not match the expected value in public databases, iSeq will attempt to redownload the files up to three times. Files that pass the MD5 check are logged as successfully downloaded, while those that fail after three attempts are recorded in a failure log.

For users requiring compressed data, iSeq can directly download FASTQ files in gzip format if available. Otherwise, iSeq will download SRA files and converts them to FASTQ format using fasterq-dump (https://github.com/ncbi/sra-tools), followed by compression with pigz (https://zlib.net/pigz), which is only considered when fetching from INSDC-related databases. For the GSA database, iSeq always directly fetches compressed or BAM format data. Additionally, iSeq can merge multiple FASTQ files from the same experiment into a single file for single-end (SE) sequencing data, or maintain the order and consistency of read names in two files for paired-end (PE) sequencing data. If the experiment consists of only one run, the run accession will be renamed in the format of the experiment accession. Finally, upon successful execution, iSeq will generate the corresponding metadata and NGS data.

## RESULTS AND DISCUSSIONS

### Design of iSeq and comparative analysis

Given the exponential growth of publicly stored NGS data and its widespread usage, we developed iSeq using Bash scripting **(Figure 2A)**. iSeq efficiently retrieves sequencing data from diverse bioinformatics repositories like GSA, SRA, ENA, and DDBJ. To simplify user retrieval processes, iSeq currently supports 25 accession formats as inputs **(Figure 2B)**, then fetches relevant metadata and sequentially downloads the included Run accessions. In fact, iSeq was inspired by existing tools like fastq-dl (https://github.com/rpetit3/fastq-dl), fetchngs (https://github.com/nf-core/fetchngs) [24], pysradb (https://github.com/saketkc/pysradb) [25], and Kingfisher (https://github.com/wwood/kingfisher-download). However, given the incomplete functionalities and database support of these tools, iSeq was devised.

**Figure 2.**
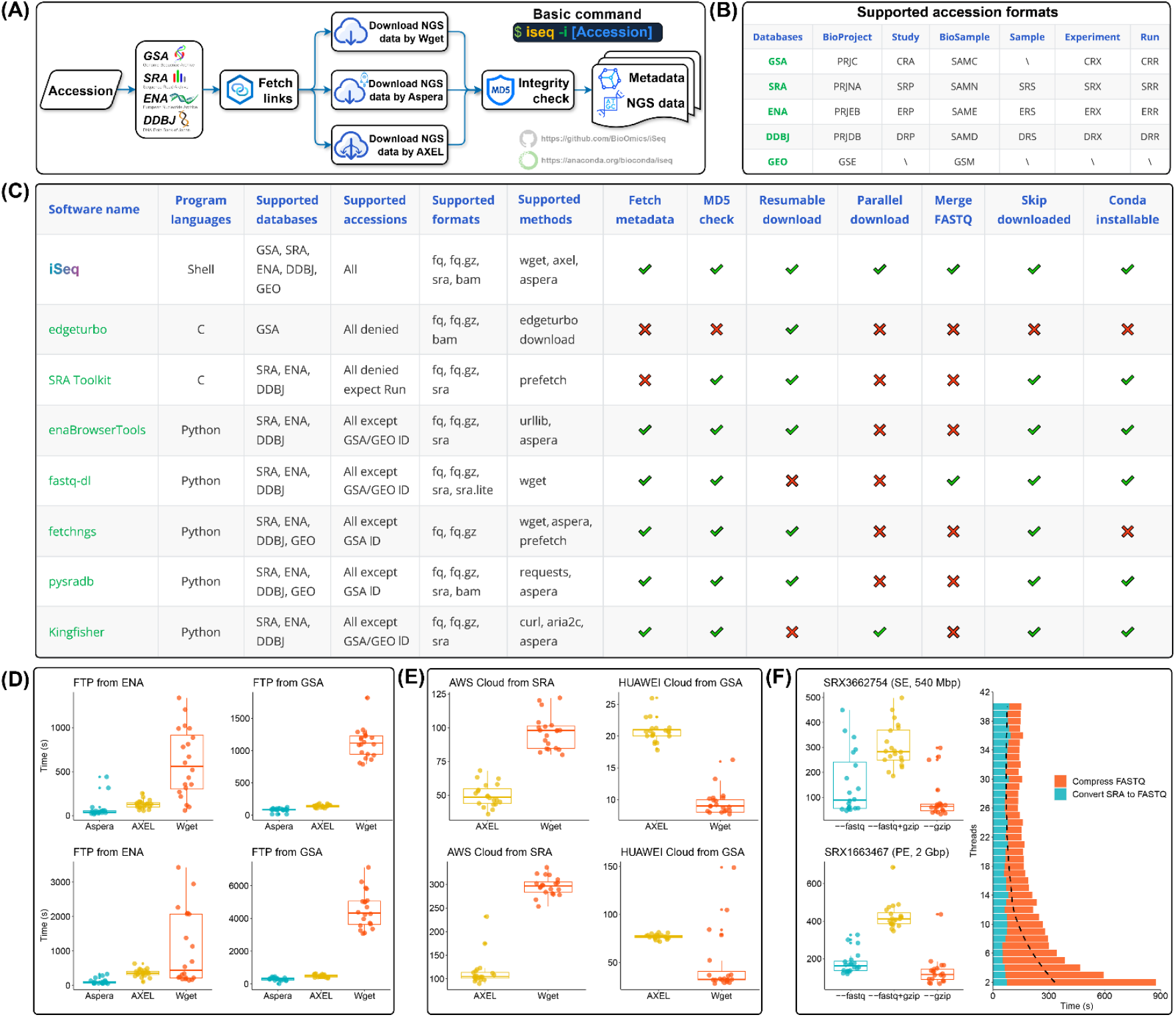
Comparison of iSeq with existing tools and performance benchmarks. **(A)** Overview of iSeq design. **(B)** Supported accession formats by iSeq. **(C)** Comparative analysis of iSeq and existing tools. **(D)** Download speed comparison using different methods. **(E)** Cloud storage download speed comparison. **(F)** Efficiency of downloading gzip-formatted FASTQ Files.

Specifically, iSeq stands as the shell command-line tool for fetching NGS data and metadata. Compared to other tools **(Figure 2C)**, it introduces support for the GSA database and its accession formats. Additionally, in terms of supported NGS data formats, iSeq allows retrieval of FASTQ, gzip-formatted FASTQ, SRA, and BAM file formats. For SRA formats, iSeq allows retrieval of SRA Normalized, the original format with full base quality scores. However, iSeq currently does not support SRA Lite, which reduces file size by simplifying quality scores to a uniform Q30. This decision is based on the potential impact of base quality scores on downstream omics analyses, where inaccurate quality scores could introduce uncontrollable biases during read quality control [26]. Therefore, iSeq refrains from incorporating SRA Lite as an output format. Moreover, recognizing the time-consuming process of single-threaded retrieval for large-scale NGS data via Wget, in addition to integrating Aspera, iSeq also supports multi-threaded downloading using AXEL for individual files. However, multi-threaded downloading may result in data format inconsistencies. Therefore, iSeq conducts integrity checks on downloaded NGS data to ensure the accuracy of downstream analysis. Additionally, in the event of unforeseen interruptions during the iSeq program process, it includes breakpoint reconnection functionality, maximizing the integrity of NGS data retrieval processes.

### Benchmarks

Firstly, to assess the stability of NGS data downloads using iSeq, we attempted to fetch 3,000 gzip-formatted FASTQ files (∼7 Terabyte of base pairs, Tbp) generated from RNA-Seq studies from the GSA database. The results showed that after the initial download, 2,998 files were successfully downloaded and passed integrity checks **(Table S1)**. Subsequently, we tried to obtain 3,000 SRA files (∼5 Tbp) used in multi-omics studies from INSDC-related databases. Ultimately, 2,999 files were successfully downloaded and passed integrity checks **(Table S1)**. Upon rerunning the same command with iSeq, it automatically skipped the already completed downloads and successfully fetched all NGS data and metadata.

Next, we evaluated the efficiency of iSeq in fetching NGS data through different channels and methods. For accessing NGS data in INSDC-related databases, iSeq prioritizes the FTP channel in ENA. We performed 20 repeated downloads of SRX3662754 (SE, 540 Mbp) and SRX1663467 (PE, 2 Gbp) using the three methods supported by iSeq. The results showed that using Aspera (Command: iseq -i [accession] -a) to fetch NGS data provided the fastest and most stable download speeds **(Figure 2D)**. Considering that Aspera requires specific firewall ports and an additional private key file (which iSeq is pre-configured with automatically when installed using conda), iSeq also supports parallel downloads using AXEL (Command: iseq -i [accession] -p [number of parallel]), achieving speeds close to Aspera **(Figure 2D)**. This conclusion holds true for the GSA FTP channel as well **(Figure 2D)**.

With the development of cloud technology, SRA and GSA have migrated data to cloud storage. Since Aspera relies on the FASP protocol, it cannot access data over the HTTPS protocol. Therefore, for cloud-stored data, iSeq exclusively uses AXEL and Wget. The results showed that AXEL, when accessing data via AWS Cloud channel from SRA (Command: iseq -i [accession] -p [number of parallel] -d sra), achieved faster download speeds **(Figure 2E)**, whereas Wget was slightly faster when accessing data via HUAWEI Cloud channel from GSA (Command: iseq -i [accession]) **(Figure 2E)**.

Lastly, we evaluated efficiency of iSeq in downloading gzip-formatted FASTQ files through different methods. The results indicated that directly fetching gzip-formatted FASTQ files (Command: iseq -i [accession] -g) was the fastest and most computationally efficient method **(Figure 2F)**. However, the “--fastq” parameter is particularly useful for downloading single-cell data, especially for scATAC-Seq data, as it can effectively decompose the files into four parts: Index, Barcode, Read 1, and Read 2. If FASTQ files are directly downloaded using the “--gzip” parameter, only Read 1 and Read 2 files are obtained (e.g., SRR13450125), which may cause issues during subsequent data analysis. Therefore, iSeq can simultaneously use the “--fastq” and “--gzip” parameters (Command: iseq -i [accession] -q -g -t [number of threads]), first fetching SRA files, then decomposing and compressing them. Notably, increasing the number of threads does not linearly reduce time. The results indicated that increasing the number of threads did not significantly enhance the speed of decomposing SRA files, but it did notably improve compression speed **(Figure 2F)**. However, when the number of threads exceeded 15, further increases in threads did not result in a substantial reduction in time **(Figure 2F)**.

In summary, users can achieve faster and more stable download speeds for FTP channel data by setting the “--aspera” parameter in iSeq, regardless of whether it’s for SRA format data or gzip-formatted files. For cloud-stored data, the same objective can be attained by setting the “--parallel [number of parallel]” parameter in iSeq. It’s important to note that actual download speeds may still vary depending on the user’s network environment and the real-time throughput of public databases.

## CONCLUSION

iSeq stands as a command-line tool, adept at accommodating diverse input formats and providing multifaceted support. In an era characterized by the continuous expansion of NGS data and the popularity of large language models, iSeq empowers researchers with the convenient and swift ability to fetch NGS data and metadata. We believe that through iSeq, the mysteries hidden within NGS data can be brought to light once more.

## Supporting information

https://github.com/BioOmics/iSeq/blob/main/docs/benchmark/iSeq-Table%20S1.xlsx

## AUTHOR CONTRIBUTIONS

**Haoyu Chao** conducted the sample collection, conceptualization, methodology, data analysis, curation, visualization, and the initial drafting and revision of the manuscript. **Zhuojin Li** debugged the iSeq program. Notably, **Ming Chen** conceived the study. **Ming Chen** and **Dijun Chen** jointly provided overall conceptualization and supervision. All authors reviewed and approved the submitted version.

## ACKNOWLEDGMENTS

The authors are grateful to the members of Ming Chen’s laboratory for helpful discussions and valuable comments.

## DATA AVAILABILITY STATEMENT

NGS data utilized in this study are publicly available from various repositories. Specifically, experiments SRX3662754 and SRX1663467 were retrieved from ENA and SRA, respectively. Additionally, experiments CRX917377 and CRX095512 were obtained from GSA. Furthermore, iSeq is accessible to all researchers via GitHub (https://github.com/BioOmics/iSeq) or Bioconda (https://anaconda.org/bioconda/iseq) [27], and users can install it on any Linux system. The code and data used for performance benchmarks can also be accessed on GitHub (https://github.com/BioOmics/iSeq/tree/main/docs/benchmark).

## SUPPORTING INFORMATION

**Table S1:** Summary of NGS data download and integrity check results.

